# Arkypallidal neurons in the external globus pallidus can mediate inhibitory control by altering competition in the striatum

**DOI:** 10.1101/2024.05.03.592321

**Authors:** Cristina Giossi, Jyotika Bahuguna, Jonathan E. Rubin, Timothy Verstynen, Catalina Vich

## Abstract

Reactive inhibitory control is crucial for survival. Traditionally, this control in mammals was attributed solely to the hyperdirect pathway, with cortical control signals flowing unidirectionally from the subthalamic nucleus (STN) to basal ganglia output regions. Yet recent findings have put this model into question, suggesting that the STN is assisted in stopping actions through ascending control signals to the striatum mediated by the external globus pallidus (GPe). Here we investigate this suggestion by harnessing a biologically-constrained spiking model of the corticobasal ganglia-thalamic (CBGT) circuit that includes pallidostriatal pathways originating from arkypallidal neurons. Through a series of experiments probing the interaction between three critical inhibitory nodes (the STN, arkypallidal cells, and indirect pathway spiny projection neurons), we find that the GPe acts as a critical mediator of both ascending and descending inhibitory signals in the CBGT circuit. In particular, pallidostriatal pathways regulate this process by weakening the direct pathway dominance of the evidence accumulation process driving decisions, which increases the relative suppressive influence of the indirect pathway on basal ganglia output. These findings delineate how pallidostriatal pathways can facilitate action cancellation by managing the bidirectional flow of information within CBGT circuits.

## Introduction

Imagine standing at the top of a ski slope, poised to descend. Just as you are about to start, something captures your attention in your periphery, prompting you to suddenly halt. A fearless skier then shoots past you, narrowly avoiding a collision. This scenario highlights how the fast suppression of a planned action in response to an external stimulus, known as *reactive inhibition* [1], can be crucial for survival.

The classical model of reactive inhibition posits that action cancellation solely depends on the hyperdirect pathway, which directly drives inhibition of the thalamus by ramping up inhibitory signals from the internal globus pallidus (GPi) via activation of the subthalamic nucleus (STN) [2]. According to this framework, the hyperdirect pathway acts as a brake, facilitating the immediate termination of upcoming actions [3, 4]. However, the dynamics of STN firing elicited by an external stop cue is largely inconsistent with this classical model, exhibiting a fast and brief burst that precedes, but is not coincidental with, basal ganglia output signals that drive suppression of the thalamus [5]. Alternatively, recent research has highlighted the involvement of a previously underappreciated cell type within the external globus pallidus (GPe), the arkypallidal neurons (Arky), that appear to be crucial for external stop cues to induce termination of a planned action [5, 6, 7, 8]. These neurons, characterized by their ascending GABAergic projections to the striatum, target both spiny projection neurons (SPNs, split into direct, dPSNs, and indirect, iSPNs, pathway neurons) and fast-spiking interneurons (FSIs) [9, 10], making the pallidostriatal pathways potential drivers of reactive inhibition by recruiting striatal systems into the cancellation process [11, 12, 13, 14, 15].

While it is now clear that pallidostriatal pathways play a role in the inhibition of planned actions, the mechanics of their influence remain unclear. To understand this, we employed a biologically-constrained spiking neural network model of cortico-basal ganglia-thalamic (CBGT) circuits, integrating recent empirical findings on the pallidostriatal pathways and simulating behavior in the stop signal task. We first replicated experimental findings on action inhibition by delivering a stop signal, separately, to each of three critical cell populations within the CBGT network (STN, Arky, or iSPN) using an external excitatory drive meant to simulate optogenetic activation. We then explored the causal role that pallidostriatal pathways play in reactive inhibition by suppressing arkypallidal activity or modifying the efficacy of pallidostriatal connections to SPNs during additional simulations of optogenetic activation. Our analysis specifically elucidates how the GPe can modulate the competition between the direct and indirect striatal pathways that mediates evidence accumulation during decision-making [16, 17, 18].

## Results

### Simulating reactive inhibition

We used our computational model of CBGT network to simulate a normative version of the well-studied stop-signal task (Figure 1A) [19, 20, 21]. Each trial begins with the presentation of an imperative stimulus (the *Go* cue), that drives cortical projections targeting the striatal SPNs. The trial decision phase reflects the entire evidence accumulation window, where iSPNs and dSPNs in the striatum compete to control the signals generated by the thalamic region targeted by CBGT outputs [16, 18, 17]. When the thalamic firing rate reaches a threshold (30 Hz), an action is triggered, and the time between the imperative stimulus and the action onset is recorded as the *reaction time* (RT). Note that our RT measure lacks the movement time component of traditional RTs recorded experimentally. At 70 ms after the imperative stimulus onset, we present the *stop signal*. This is represented as an injection of a boxcar-shaped excitatory current into a target cell population. This current boosts the activity of the targeted cells, with the goal of reducing the likelihood that the thalamic activity will reach the action threshold. An action is considered to be successfully stopped if the thalamic threshold is not reached within the trial window (300 ms) and therefore no action is registered.

**Figure 1:**
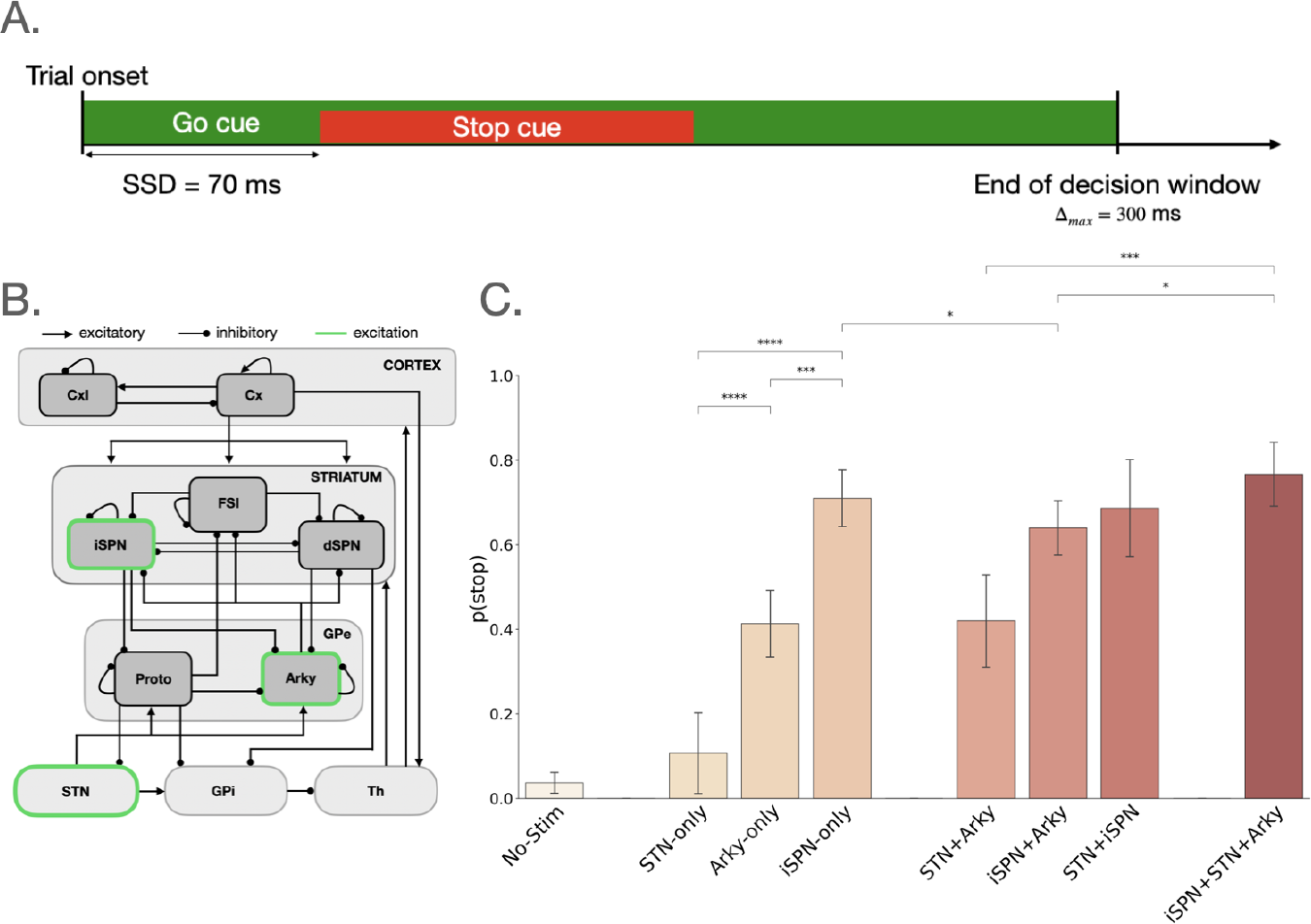
Simulating reactive inhibition. (A) Schematics of our computational implementation of the stop signal task paradigm. The trial onset corresponds to the presentation of a primary stimulus (the “Go” cue, in green), which is sustained until the end of the decision window. A secondary stop stimulus follows (“Stop” cue, in red), instructing the network agent to withhold the action selection process. The delay between the onsets of the two stimuli is referred to as stop signal delay (SSD; 70 ms). The maximum duration of the decision phase corresponds to Δ_*max*_ = 300 ms. (B) CBGT network architecture. Arrows depict excitatory projections while circles represent inhibitory connections. Green outlined nuclei indicate the populations that were externally stimulated to simulate the effects of the stop signal presentation. Cx: cortex; CxI: cortical interneurons; FSI: fast-spiking interneurons; iSPN: indirect pathway spiny projection neurons; dSPN: direct pathway spiny projection neurons; GPe: external globus pallidus; Proto: GPe prototypical neurons; Arky: GPe arkypallidal neurons; STN: subthalamic nucleus; GPi: internal globus pallidus; Th: thalamus. (C) Stopping probability across different stimulation cases. Individually, iSPN stimulation produces a ∼ 71% of chance of stopping, Arky ∼ 41%, and STN ∼ 11%. The simultaneous stimulation of all three nuclei produces an overall greater likelihood of stopping (∼ 77% stopping probability). t-test paired samples with Bonferroni correction; p-value annotation legend: *: 0.01<p<=0.05; **: 0.001<p<=0.01; ***: 0.0001<p<= 0.001; ****: p<=0.0001

For this study, we injected the stop signal into one of three CBGT cell populations that have been experimentally shown to inhibit behavior: iSPNs [22], STN [4], and Arky [6, 5] (Figure 1B). The iSPNs are the primary input of the indirect pathway and are traditionally considered to drive proactive inhibitory control by suppressing the thalamus as part of the direct/indirect pathway competition [23, 24, 16]. The STN cells, on the other hand, reflect the primary basal ganglia input from the cortical hyperdirect pathway [2]. This circuit bypasses the striatum altogether and is thought to provide a fast control mechanism for action suppression [3]. Finally, the recently rediscovered Arky cells [25] provide inhibitory projections to striatal iSPNs, dSPNs, and FSIs [9, 26, 27, 28, 29, 10, 30, 7, 6, 31]. Importantly, activation of Arky cells has recently been shown to suppress actions [5, 8, 6].

Activation of these target populations individually achieved varying degrees of action suppression (ANOVA test across stimulation conditions: F[7,29]=113.16, p<0.0001; Figure 1C). Stimulating the STN alone had the weakest influence on stopping, with an 11% stopping probability that improved only marginally over the 4% chance ”stopping” probability during trials when no stop signal was delivered. These probabilities were far lower than the stopping probabilities that we observed with Arky-only (41%) and iSPN-only (71%) stimulation. We next considered the stimulation of pairs of regions, with the best performance, a 69% stopping probability, occurring with simultaneous STN and iSPN stimulation, comparable to the iSPN-alone condition. Finally, when all three regions were stimulated simultaneously, we observed a 77% stopping probability, which is slightly higher than the iSPN-only stimulation. These results show that, as nodes higher up in the standard basal ganglia hierarchy are targeted, the efficiency of action cancellation increases. While aspects of the action suppression resulting from STN-driven inhibitory control (via direct excitation of the internal globus pallidus; GPi) and iSPN-driven control (via the long and short indirect pathways) are well understood, the mechanism by which Arky cells drive, or otherwise contribute, to this behavioral inhibition remains unclear. We next explore the contributions of this population more deeply.

### Impact of arkypallidal neurons on reactive inhibition

To analyze how Arky neurons impact reactive inhibition, we measured how their activation alters the overall dynamics of the CBGT circuit, particularly its postsynaptic targets in the striatum. Stimulating Arky cells pushed the rest of the CBGT network into an overall movement-suppressive state (Figure 2A). Notably, the dSPNs exhibited the strongest drop in firing rates due to Arky activation, followed by the iSPNs. Downstream, we observe that these changes result in an increase in GPi activity and, consequently, decreased activity in the thalamic targets of the basal ganglia.

**Figure 2:**
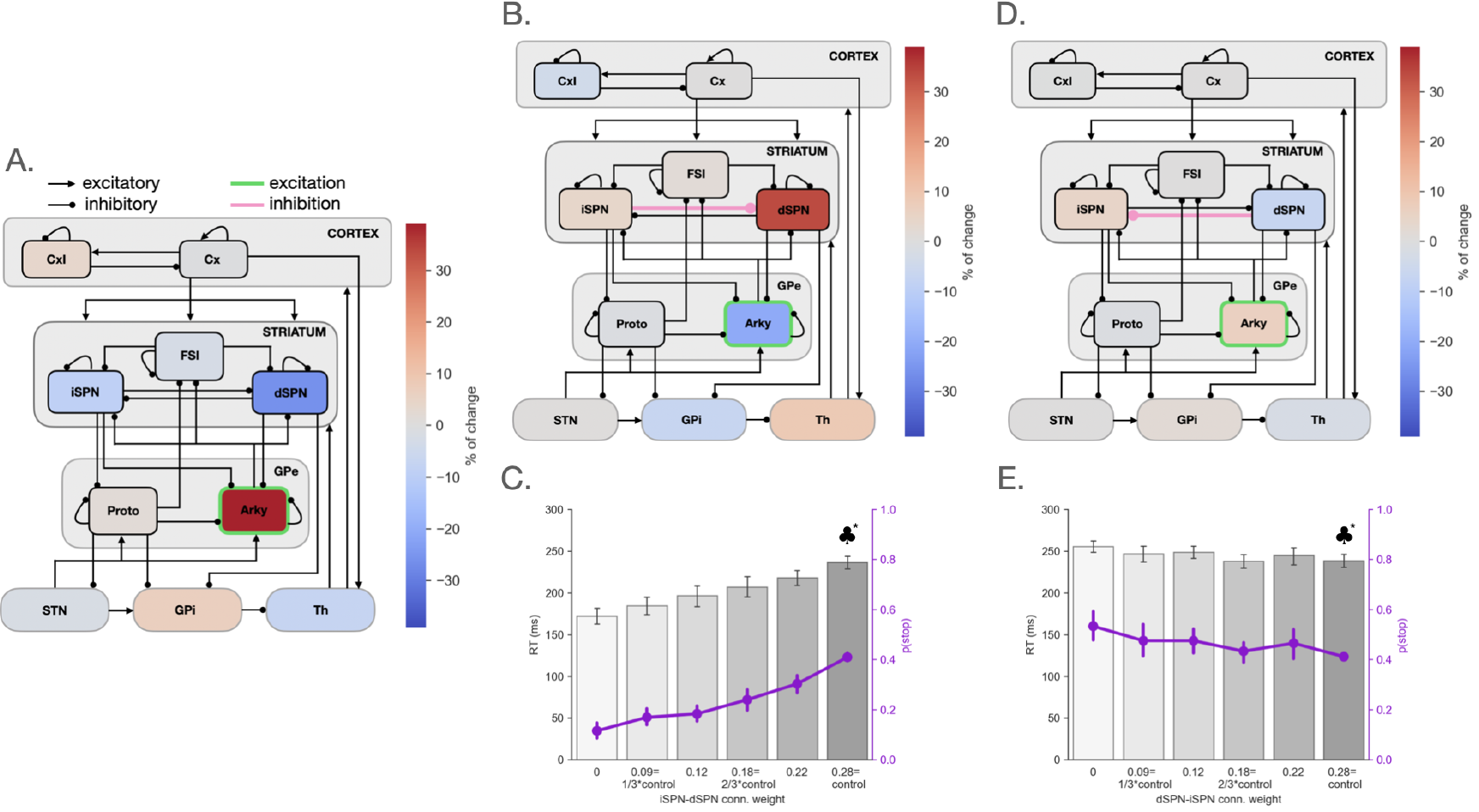
Stimulation of Arky cells shifts the balance between striatal populations. (A) Stimulation of Arky neurons (green outline) induces a stronger decrease in dSPN firing than in iSPNs. (B), (D) The change in the CBGT network’s firing patterns during Arky stimulation (green), while the reciprocal collaterals between iSPNs and dSPNs are individually attenuated (pink connections). Firing rate changes show the change between the two cases of extreme collateral connection strengths, 0.28 (control) and 0. (C), (E) Changes in network’s ability to suppress responses and in RTs when the experiments depicted in panels B and D, respectively, are implemented in steps. ^*^♣ depicts baseline case.

This asymmetric influence of Arky stimulation on i/dSPN activity suggests that Arky neurons influence basal ganglia output by affecting the competitive balance between the direct and indirect pathways [16, 18, 17]. To explore this idea further, we stimulated Arky cells while altering the strength of the reciprocal inhibitory collaterals between iSPNs and dSPNs. Gradually removing the inhibition provided by iSPNs to dSPNs during Arky stimulation produced an increase in dSPN activity that, in turn, resulted in a reduction of the network’s ability to withhold a decision (F[5,29]=36.25, p<0.0001; Figure 2B-C) and produced faster reaction times on failed stop trials (F[5,29]=17.42, p<0.0001; Figure 2C). At first glance, a reduction in Arky cell activity during stimulation might seem counterintuitive. However, this effect is attributed to a recurrent loop between Arky cells and dSPNs: as dSPN firing intensifies, due to the release from inhibition from its iSPN afferents, dSPN subsequently exerts greater inhibition on Arky neurons, leading to a decrease in their activity. When we tested the reverse scenario, by gradually attenuating the inhibition exerted by dSPNs onto iSPNs during stimulation of the Arky population, we observed a subsequent rise in the activity of iSPNs. This change led to a marginal enhancement in the probability of successful stopping (F[5, 29]=2.44, p=0.046; Figure 2D). Additionally, this also resulted in slightly longer reaction times (F[5,29]=2.18, p=0.07; Figure 2E). These results confirm that the suppression of actions induced by activation of Arky cells is moderated by a shift in the balance of power against the direct and in favor of the indirect pathway.

Because they receive input from the STN and convey inhibitory output to the striatum, the Arky neurons are ideally positioned to be a central node in an ascending pathway that can modulate the descending control signals from the striatum that drive reactive inhibition in CBGT circuits [11]. To further evaluate their influence, we simulated a lesion of Arky neurons by injecting a suppressive current into these cells while a stop signal was delivered to the STN cells, iSPNs, or both. In each experiment we compared network responses with Arky cell suppression (lesion) to conditions where the Arky cells were left unperturbed (control).

We started with the combined stimulation case (i.e., stimulation of both STN cells and iSPNs; Figure 3A), where we observed a discernible decline in the likelihood of successful stops when the Arky cells were suppressed (control: 0.74 ± 0.0116 (± SEM), lesion: 0.60 ± 0.0268; t(9)=4.44, p=0.0016; Figure 3B). This effect was not reliably observed in the RTs (control: 285 ± 4, lesion: 282 ± 2; t(9)=0.94, p=0.3721; Figure 3B). We next aimed to dissect the dynamics induced by the simultaneous stimulation of the two nuclei by independently activating each of them. We first looked at the involvement of Arky cells in the transmission of descending indirect pathway control signals originating from the iSPNs (Figure 3C). Here we observed similar effects as in the combined stimulation case, with a noticeable decrease in the likelihood of successful stops (control: 0.70±0.0060; lesion: 0.53 ± 0.0219; t(9)=6.99, p<0.0001; Figure 3D). As before, a change was not consistently observed in RTs (control: 282 ± 3, lesion: 276 ± 4; t(9)=2.10, p=0.065; Figure 3D). These results indicate two things. First, Arky cells play a role in modulating the effectiveness of descending control signals within the indirect pathway. Second, iSPNs play a dominant role in the effects observed from stimulating both STN neurons and iSPNs (Figure 3A-B). We next tested the classical stop signal control model by activating the hyperdirect pathway via stimulation of the STN cells (Figure 3E). Here suppression of Arky neurons resulted in both a reduction of the network’s ability to stop effectively (control: 0.10±0.0086, mean ± SEM; lesion: 0.01 ± 0.0075; t(9)=6.23, p=0.0002; Figure 3F) and the RTs observed on trials where stopping did not occur (control: 180 ± 3; lesion: 139 ± 2; t(9)=9.06, p<0.0001; Figure 3F). This outcome confirms that Arky cells also influence the effectiveness of hyperdirect pathway control, including RTs. Although the hyperdirect pathway is traditionally viewed as a descending pathway, from STN to basal ganglia output nuclei, these results also indicate that their ascending component, gated by Arky cells that project to the striatum, significantly affects the network’s capacity to stop or slow actions.

**Figure 3:**
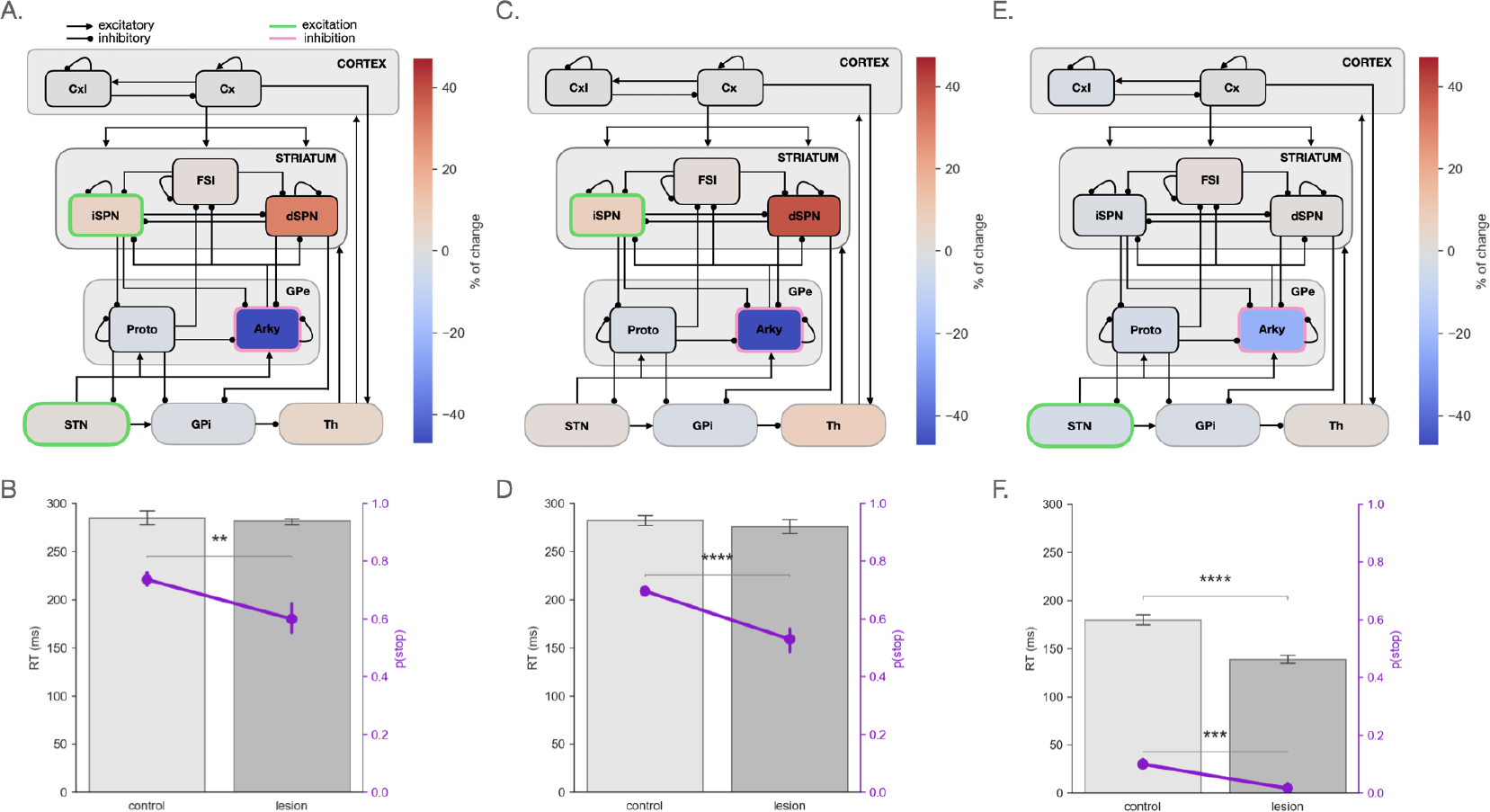
Arky cells mediate ascending and descending information flow within the CBGT network in the context of reactive inhibition. (A), (C), (E) Network schematics of experiments where Arky population activity is suppressed while the STN and iSPNs (A), iSPNs alone (C), or STN alone is stimulated (E). Color codes show the changes in firing rates after suppression of the Arky neurons compared to before. (B), (D), (F) Stopping probability and reaction time (RT) across the same task conditions portrayed in panels A, C, and E. Lesion refers to the case with suppression of Arky cell activity.

These findings suggest that the pallidostriatal pathways contribute to the bidirectional flow of inhibitory control signals within the CBGT circuit. To see how Arky cell suppression impacts the activity of the network, we also calculated the firing rate changes from before to after Arky suppression in each of the three conditions shown in Figure 3. With the stop signal being delivered only to the STN neurons, Arky cell suppression had a relatively small impact on overall network activity (node colors in Figure 3E). In contrast, for both iSPN stimulation conditions (3A-C), suppressing Arky cells led to amplified activity in both SPN populations, with a stronger effect on dSPNs than iSPNs. These observations suggest that Arky neurons modulate the dynamics downstream of the striatum primarily by helping to suppress dSPN activity.

### Mechanisms underlying Arky effects on reactive inhibition

The preceding experiments show how pallidostriatal pathways likely influence striatal activity, and hence descending information flow, by shifting the relative dominance away from dSPNs and toward iSPNs. To directly test this idea, we drove reactive inhibition by delivering the stop signal to both iSPNs and Arky cells while separately changing the efficacies of the pallidostriatal connections to the iSPNs and the dSPNs. As we reduced the strength of Arky GABAergic projections onto iSPNs, we observed an increase in iSPN activity. This, in turn, contributed to a further decrease in dSPN firing rates, as well as a drop in Proto cell activity (Figure 4A). These changes produced an increase in GPi firing, subsequently leading to a reduction in thalamic activity. These effects enhanced the network’s ability to suppress responses (F[4,29]=4.23, p=0.005; Figure 4B), accompanied by longer RTs (F[4,29]=4.31, p=0.005; Figure 4B). Conversely, as we gradually reduced the Arky inhibition to dSPNs, we observed a dramatic increase in dSPN activity, which lowered the firing rates of both iSPNs and Arky neurons (Figure 4C; note the change in scale of effects between Figures 4A and 4C). This rise in dSPN activity resulted in decreased GPi activity, which in turn led to an amplification of thalamic firing (Figure 4C). Consequently, the network failed to adequately suppress responses (F[4,29]=132.28, p<0.0001; Figure 4D) and produced shorter RTs on missed stop trials (F[4,29]=50.22, p<0.0001; Figure 4D). Together with Figure 2A, these results suggest that Arky neuron inputs to dSPNs and iSPNs are instrumental in modulating the balance of power between these striatal populations, such that the bias of the network shifts to the indirect pathway as Arky cells increase their activity.

**Figure 4:**
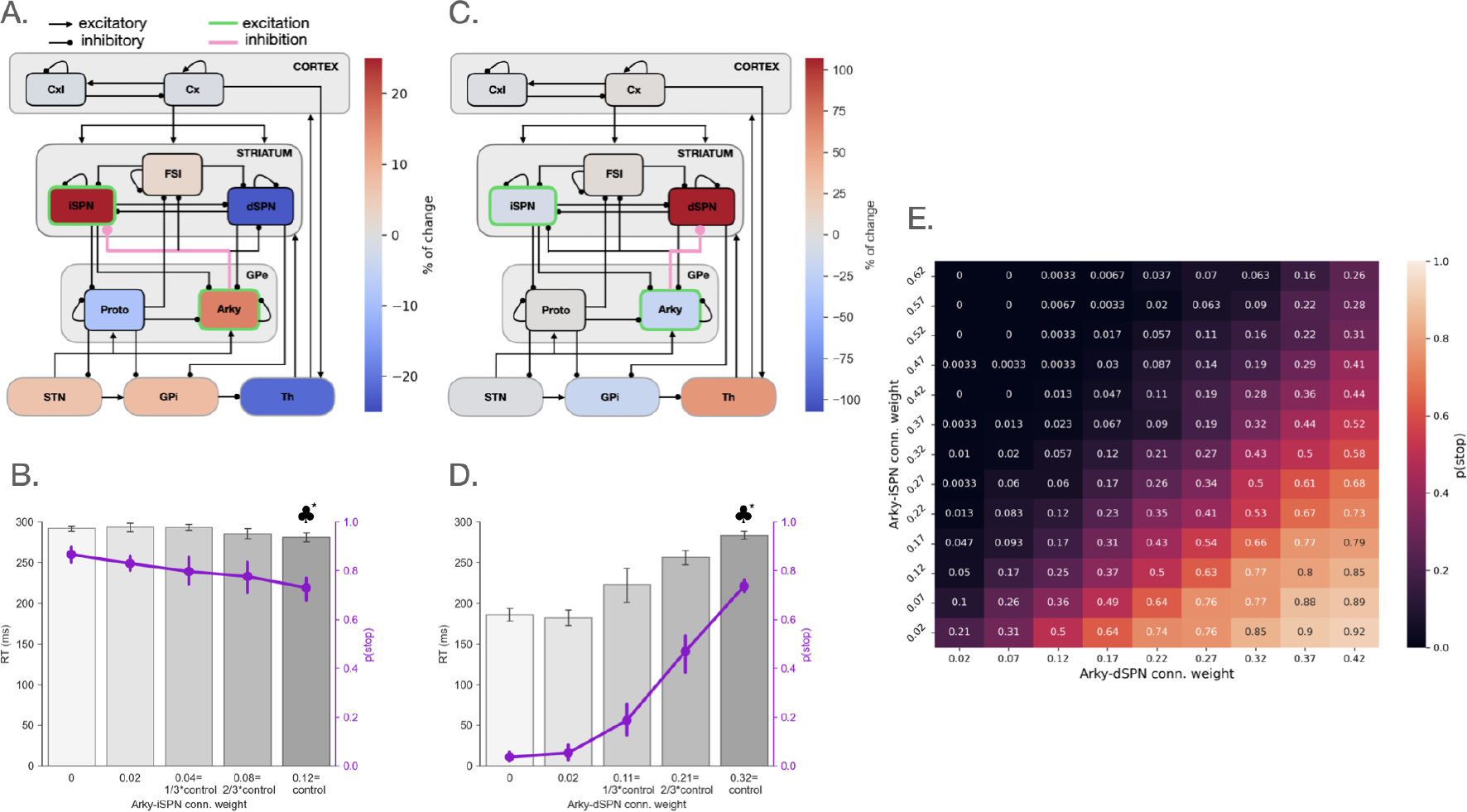
Arky neurons shift the striatal balance of power due to differential inhibition of dSPNs and iSPNs. (A), (C) Network firing schematics during Arky neuron and iSPN stimulation, with concurrent blocking of pallidal inhibitory input into iSPNs or dSPNs, respectively (in pink). Firing rate changes show the change between the two extreme conditions of 0 and 0.12 (control) for Arky → iSPN manipulation (panel A) and 0 and 0.32 (control) for Arky → dSPN manipulation (panel C). (B), (D) Evolution of network’s ability to suppress responses and of RTs during the experiments depicted in panels B and D, respectively. ^*^♣ depicts for baseline behavior. (E) Synaptic weight analysis to study the relative strengths of inhibitory projections from Arky cells to striatal populations. The effect of iSPN and Arky population stimulation grows with the ratio of Arky → dSPN to Arky → iSPN connection strengths.

To provide a more thorough test of our hypothesis that the Arky cells tune the competition between direct and indirect pathways, we conducted a synaptic weight analysis, varying the relative degrees of Arky → iSPN and Arky → dSPN inhibition, while delivering a stop signal to both iSPNs and Arky cell populations. As expected, boosting the relative inhibition of dSPNs over iSPNs increased the overall stopping probability (Figure 4E), consistent with our hypothesis.

## Discussion

Recent empirical observations have emphasized the contribution of Arky neurons in the rodent GPe for promoting action suppression [5, 8, 6, 7]. Yet how this control is implemented remains unclear. We constructed a biologically-constrained spiking neural network model of the CBGT circuitry, using the avialable empirical evidence to define the connectivity and physiology of the model, to simulate performance in a normative version of the stop signal task. By reproducing the effects of activating three major stop-related targets within the CBGT circuit, we found that nodes higher in the CBGT hierarchy exhibited greater influence on action cancellation. Our results place pallidostriatal pathways as a central hub of the CBGT network, regulating the back-and-forth transfer of signals within the basal ganglia circuitry. In particular, our predictions elucidate how pallidostriatal pathways tune reactive control by altering the competition between direct and indirect pathways via decreases in the relative influence of the direct pathway.

Our simulation results align very closely with prior empirical observations. Schmidt et al. (2013) [32] discovered that the cortical-STN-SNr (hyperdirect) pathway reacts swiftly to stop cues, well in advance of the stop signal reaction time. Their results suggest that this mechanism may be too rapid to be fully responsible for preventing actions. Moreover, it may be insufficiently specific, as similar responses have also been observed for go cues in similar tasks [33, 5]. Building upon this evidence, Mallet et al. (2016) [5] proposed a two-step model for reactive stopping. In this model, the hyperdirect pathway initiates a rapid ”Pause” signal, allowing time for a slower, separate ”Cancel” process to occur within the striatum. Notably, an increase in firing rates has been observed in both Proto and Arky neurons following the presentation of a stop signal, with Arky neurons showing a more pronounced response than Proto cells, consistent with their involvement in halting actions. This observation was confirmed by Aristieta et al. (2021) [6], who achieved abrupt interruption of actions by optogenetically stimulating Arky neurons in rodents engaged in treadmill locomotion, an effect that the authors suggest is likely mediated through suppression of striatal activity. Pamukcu et al. (2020) [7] demonstrated that optogenetic stimulation of the axon terminals in the dorsal striatum from Npas1-expressing neurons in GPe, 60% of which are Arky neurons, decreased motor output. Collectively, these experimental findings underscore the pivotal contribution of the pallidostriatal pathways in inhibitory control. Importantly, our simulations build on these results by providing evidence that without Arky neuron involvement, the slowed RTs associated with STN stimulation would be lost, and the probability of stopping under iSPN stimulation would be reduced. Thus, Arky neurons appear to play a critical role in integrating control signals from both hyperdirect and top-down pathways.

Emerging evidence will likely continue to update our understanding of how Arky cells centrally mediate the ascending and descending flow of control signals in CBGT pathways. For example, there is some evidence to suggest that the Arky cells receive their own direct inputs from cortex [34]. Such inputs would extend the role of GPe beyond simple mediation of hyperdirect and direct/indirect pathway control signals. Other evidence contradicts this idea of direct cortical control of Arky cells, however, indicating no specific bias in cortical projections towards any GPe neuron subtype [35, 36]. In addition, there may be monosynaptic connections from STN to striatum that facilitate the transmission of hyperdirect signals. The influence of these connections on the activity of SPNs remains uncertain, although there appears to be a significant impact on striatal parvalbumin (PV) interneurons [37]. These and other emerging details may expand our view of the mechanisms of pallidostriatal control but will not change the fact that palli-dostriatal pathways mediate ascending and descending control signals.

Using optogenetics to replicate our simulation and lesion experiments *in vivo* could provide critical tests of our model predictions and hence deeper insights into the contributions of CBGT pathways to reactive inhibition. Fiber photometry could yield valuable information about the uniformity of Arky cell involvement in various inhibitory control pathways, including whether inputs from various nuclei target distinct Arky neuron subgroups. In light of such complexities, it becomes imperative to dig deeper into the dynamics of inhibitory control within CBGT circuits and elucidate the nuanced contributions of the pallidostriatal pathways. Such investigations will be crucial for achieving a comprehensive understanding of how CBGT circuits shape behavioral control.

## Materials and Methods

### CBGT network

The CBGT model is a computational, biologically-inspired spiking neuronal network, comprising six distinct neural regions: the cortex, with excitatory (Cx) and inhibitory (CxI) subpopulations; the striatum, including dSPNs, iSPNs, and FSIs; the GPe, segmented into Proto and Arky subpopulations; the STN; the GPi; and a thalamic component (Th). Figure 1B illustrates the network’s connectivity, where three main pathways can be distinguished. In the *direct pathway*, cortical inputs activate the dSPNs, which in turn inhibit the GPi, causing disinhibition of thalamic activity, potentially facilitating action selection. Conversely, the *indirect pathway* involves cortical inputs activating the iSPNs, which inhibit the GPe, forming the so-called ”short” (Proto-GPi) and ”long” (Proto-STN-GPi) indirect pathway. By optogenetically stimulating the STN, we also simulate the activation of the *hyperdirect pathway*. This third pathway is known to operate independently of the striatum, directly regulating thalamic inhibition through cortical input to the STN and its excitatory influence on GPi. For more details on the network implementation, see [38]; see also [11] for a review of experimental findings that justify the network structure used.

### Stop signal task

We implemented a computational version of a standard stop signal task (Figure 1A), where the network must control the execution or suppression of an action, following the onset of imperative cues. The first cue presented corresponds to a ”Go” stimulus, applied to the Cx, which drives the network towards a decision. During a trial, a decision is made when the thalamic firing rate reaches 30 Hz. If no decision is made within a trial window of 300 ms, then no choice is recorded and a successful inhibition occurs. 70 ms after the Go stimulus, another cue is presented: the ”Stop” signal. We modeled this signal as a boxcar-shaped external excitatory current that we inject directly into a specific nucleus. In this computational study, we apply a stop signal current to distinct target populations implicated in the process of action suppression. For more details, see Section *Simulating reactive inhibition*. This current amplifies the activity of the targeted region. For a comprehensive understanding of the task, refer to our methods paper [38]. The stop signal current is characterized by several key parameters: (a) *amplitude*, indicating the intensity of the applied stimulation; (b) *population*, specifying the targeted CBGT region; (c) *onset*, indicating the timing of the stimulation relative to the trial onset; (d) *duration*, defining how long the stimulation lasts. The choice of these parameters influences the resulting stopping probability and reaction time distribution. For this study, we set the amplitude, onset, and duration parameters to establish a baseline condition with a stopping probability of approximately 75% (*amplitude*=0.4 Hz, *onset* =70 ms, and *duration*=145 ms). The stopping probability was calculated by averaging the frequency of instances where no decisions were made across 10 threads, each consisting of 30 trials, resulting in a total of 300 trials.

## Data sharing

The network codebase utilized in this study can be found on our GitHub repository and accessed at https://github.com/CoAxLab/CBGTPy/blob/main. Detailed installation instructions and a comprehensive list of implemented functions can be found in the README.txt file within the repository. All datasets generated and analyzed during the course of this research, along with a demonstration demo, are openly available on GitHub at https://github.com/giossic/arky-stopsignal. Any additional data and code supporting the findings of this study are available from the corresponding author upon reasonable request.

## Acknowledgements

CG and CV are supported by the PCI2020-112026 and PCI2023-145982-2 projects, both funded by MCIN/AEI/10.13039/501100011033 and by the European Union ”NextGenerationEU”/PRTR as part of the CRCNS program. CG is also supported by the Conselleria de Fons Europeus, Universitat i Cultura del Govern de les Illes Balears under grant FPU2023-008-B. JB is supported by ANR-CPJ-2023. TV, JB and JER are partly supported by NIH awards R01DA053014 and R01DA059993 as part of the CR-CNS program. JER is partly supported by NIH award R01NS125814, also part of the CRCNS program.

